# Image-based parametric finite element modelling for studying contact mechanics in human knee joints

**DOI:** 10.1101/2023.09.07.556747

**Authors:** R. Readioff, R. Seil, C. Mouton, L. Marks, O. Barrera

## Abstract

**Purpose:** This study presents a framework for generating patient-specific finite element models, parameterised and optimised for contact mechanics from computed tomography (CT) scans, by avoiding the segmentation step usually employed to transform medical images into 3D models. Two morphological parameters affecting contact mechanics were investigated in the framework development: tibial cartilage thickness and tibial spine height. This study explores the effect of the interplay of these parameters in load sharing between meniscus and articulating cartilage, meniscal posterior and anterior roots strain and menisci kinematics.

**Methods:** Morphological measurements from four knee CT scans were collected, such as the maximum thickness of the tibial cartilage (ranging from 1.1 to 5.2 mm), the height of the tibial spine (ranging from 3.55 to 10.1 mm), and the width of the tibial plateau in both the coronal (ranging from 27.3 to 36.17 mm) and sagittal (ranging from 31.79 to 53.77 mm) planes. These measurements were taken for the lateral tibial plateau for both left and right knees. Subsequently, three finite element (FE) models were generated, comprising lateral tibial plateaus, lateral femoral condyle and lateral meniscus. The tibial cartilage thickness was kept at a constant value of 1 mm while varying the tibial spine height within the range measured from the CT images. This resulted in three FE models with varying spine heights, categorised as large (height = 7.42 mm), medium (height = 4.25 mm), and small (height = 1.63 mm) tibial spine heights. The menisci in the FE models were generated to be congruent with the tibial plateau. For the first time, this study advances the representations of the knee menisci microstructure in FE modelling, such that we have generated meniscus FE models with three layers of a hyperelastic model in which layer thickness and layer-specific hyperelastic material parameters are derived from our previous experimental work.

**Results:** The load sharing between the meniscus and articular cartilage was not sensitive to the varying tibial spine heights. In all three FE models, cartilage carried more than 90% of the applied load. However, the meniscus kinematics and root strains varied considerably with changing tibial spine heights. The small tibial spine height model predicted the highest meniscus movements (8.12 and 9.33 mm in the radial and circumferential directions, respectively) and the highest root strain (21.92 and 22.19 mm/mm in the anterior and posterior roots, respectively).

**Conclusion:** Our framework can generate finite element models of patients’ knees using clinical data (i.e., CT scans) without the need for lengthy image segmentation. This process is not only time-efficient but also independent of imaging operators. The models converge quickly (¿30 minutes on 2 cores) using an implicit solver with non-linear geometry and have the capability to predict contact mechanics between the articulating surfaces, meniscus kinematics and root strains. The modelling strategy presented here can provide valuable insights into predicting changes in the mechanics of soft tissues in the knee joint. It is particularly useful for investigating injury and surgical mechanisms related to the meniscus.

## 1. Introduction

The contact mechanics of soft tissues in human knees, such as articular cartilage and menisci, play a crucial role in joint function and overall knee health [1, 2, 3, 4]. Numerical modelling techniques, such as finite element analysis, are often employed to study the contact mechanics of soft tissues in human knees. These models simulate the interaction between the soft tissues, joint geometry, loading conditions, and material properties to analyse contact pressures, stress distribution, and deformation patterns. This knowledge will provide insights into injury mechanisms, hence facilitating the development of strategies for delaying, preventing, or treating knee injuries [5, 6, 7, 8, 9, 10].

Finite element (FE) modelling of human knees is a computational technique used to simulate the behaviour and response of the knee joint under various loading conditions. Patient-specific human knee models are tailored to individual patients using their medical imaging data, such as magnetic resonance imaging (MRI) or computed tomography (CT) scans. These models aim to capture the unique anatomical and biomechanical characteristics of an individual’s knee joint. They are valuable tools in orthopaedic research, clinical decision-making, and surgical planning [11, 12, 13, 14, 15, 16]. However, developing patient-specific finite element models of the knee can be challenging and may require careful consideration to achieve numerical convergence. Various factors make these models complex, but one of the most significant is accurately capturing and meshing the intricate geometry of the knee. Generating a high-quality mesh that captures important anatomical features while maintaining computational efficiency is crucial for successful and reliable simulations. The geometries of knee joint components are commonly reconstructed from medical images (i.e., CT and MRI) using an image segmentation process in a specialised imaging software (e.g., Simpleware ScanIP, Materialise MIMICS and others). This step involves separating the bones, articulating cartilage, ligaments, and menisci from each other and creating three-dimensional (3D) geometry for each component. The segmentation process carries limitations which can affect mesh accuracy, subject-specific representations of the knee and FE analysis [17, 18, 19, 20]. For example, the segmentation process depends on the quality of the medical images and the operator’s segmentation skills in identifying the anatomy and producing congruent and smooth articulating surfaces. Unfortunately, the current imaging technology employed in routine clinical diagnosis is inadequate in generating high-quality images for producing accurate and efficient FE models of the knee, particularly the menisci and ligaments. Achieving numerical convergence can be challenging using FE models derived from image segmentation processes. Halonen et al. [21] used a patient-specific FE model generated from MRI to study cartilage strain and meniscus movement of human knees during standing. They reported convergence issues, such that their FE simulation failed to converge when applying 880 N; therefore, their models were simulated under a reduced load of 50 N. This further emphasises the limitations of employing the image segmentation process for generating FE models.

Knee arthrograms, orthoscans with a contrast agent, offer a low-cost option for directly visualising the soft tissues within the joint, including cartilage, meniscus, and ligaments. Here, we proposed a methodology to create a parametric finite element model of the human knee using arthrograms to determine knee parameters, generating congruent and simplified 3D knee models resulting in rapid numerical convergence. Our methodology involves measuring and parameterising the tibial spine height, as well as the thickness of the cartilage and menisci. These measurements can be used to create simplified 3D FE models, starting from the articular cartilage profile, which can then be used to construct a meniscus structure conformed to the patient’s characteristics. This can result in congruent articulating surfaces with a high-quality mesh, ideal for rapid numerical simulations. This paper aims to illustrate the simple and fast computational framework for creating parameterised human knee joints that can lend themselves to automation and the rapid generation of similar models for stochastic studies. Here, we aim to employ the models generated from the computational framework to investigate the effect of intra-articular parameters on the load distribution between cartilage and meniscus and meniscus kinematics.

## 2. Material and Methods

### 2.1. Morphological parameters

Four human knee arthrograms were used to extract morphological measurements of the tibia and the articular cartilage. Geometrical parameters from the lateral compartment of the knee joints, such as maximum articular cartilage thickness, maximum tibial spine height relative to the tibial plateau surface (Fig. 1a and b) and the tibial plateau width in the coronal (Fig. 1a) and sagittal (Fig. 1b) planes, were collected to guide the generation of the computational models. The maximum articular cartilage thickness was extracted by observation at the most concave area of the lateral tibial plateau from the coronal plane. The estimated parameters are recorded in Table 1.

**Table 1:**
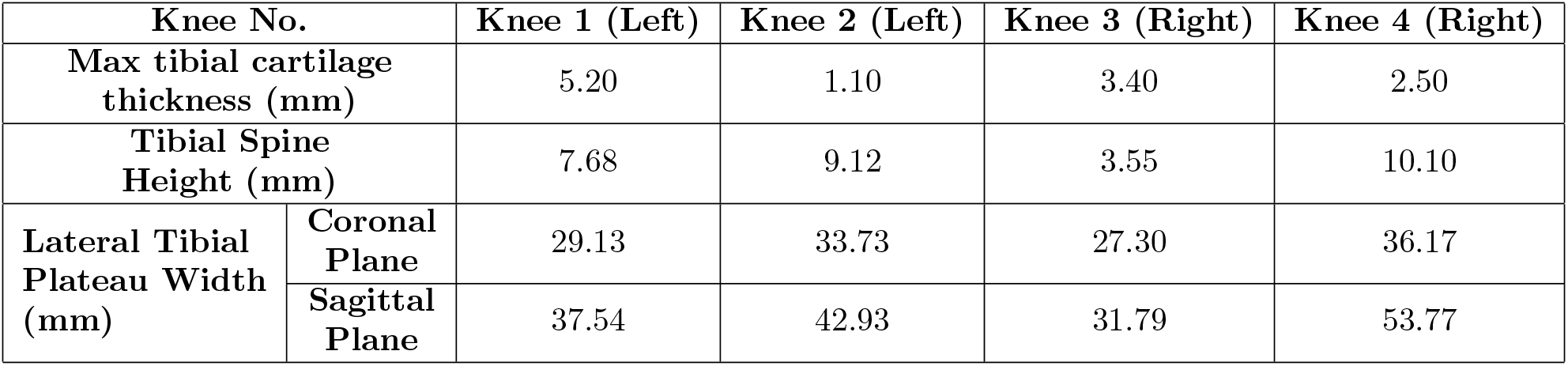
Geometrical parameters of knee joint lateral tibial plateau collected from arthrograms.

**Figure 1:**
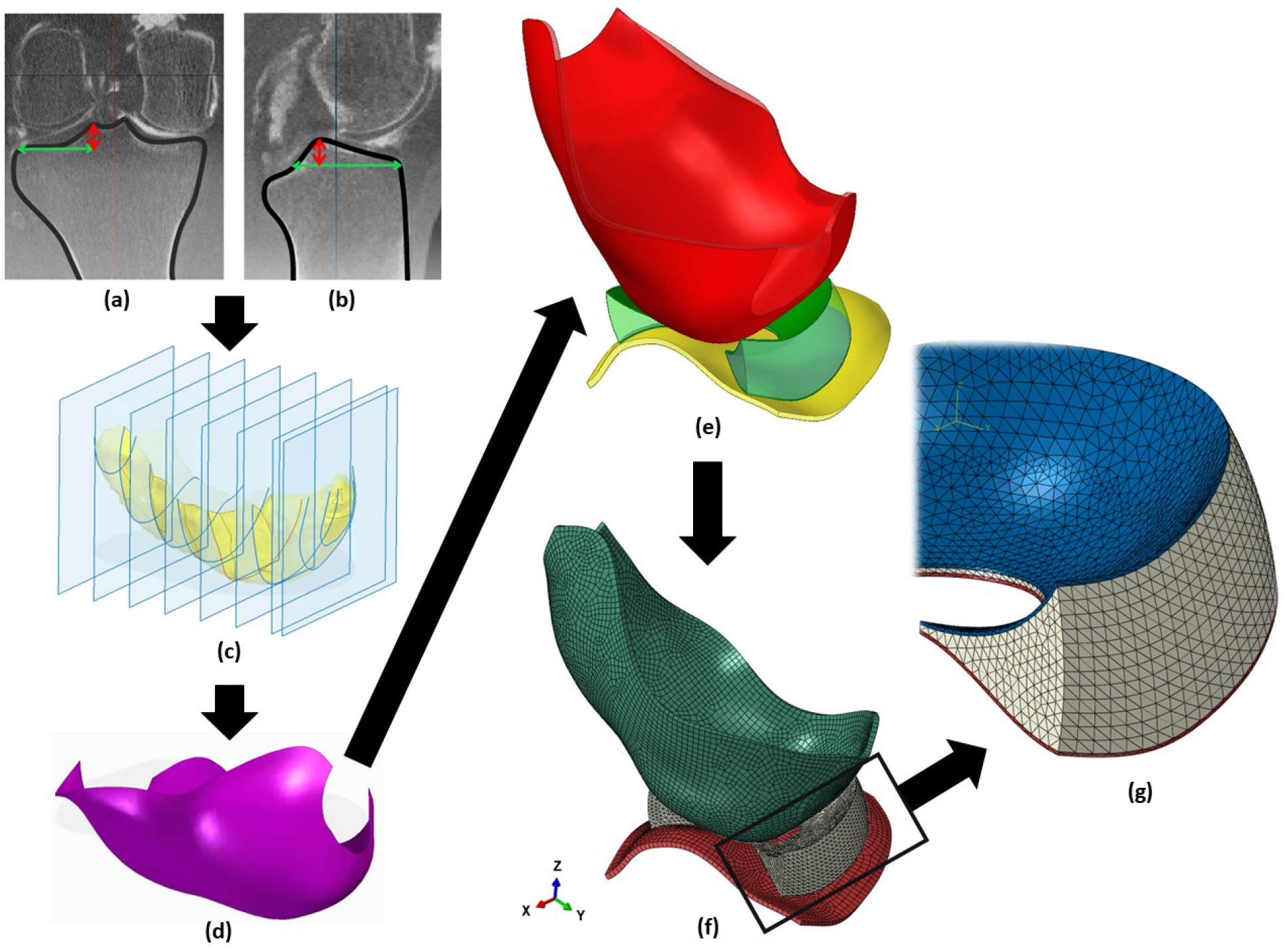
Workflow describing the generation of an adaptable (parameterised) three-dimensional (3D) shape of a lateral compartment of the knee joint, comprising the representations of tibial and femoral articular cartilage and meniscus. The parameters of the tibial plateau relative to the tibial spine from the (a) coronal and (b) sagittal arthrograms images were used in deriving (c) spline lines and generating (d) 3D surfaces of the knee components. The 3D surfaces were converted into (e) solid structures generating (f) continuous solid finite element models of the knee joint components. The meniscus component comprised (g) three layers: two thin outer layers in red and blue and the middle layer in grey.

### 2.2. Parameterised 3D geometry of lateral compartment of knee joints

The methodology for constructing parametric models of tibial and femoral cartilage, as well as meniscus geometry, has been executed using SOLIDWORKS 2021 (Dassault Systèmes). The parametric features allowed for a semi-automated process in building different geometrical models of the knee joint tibial plateau with varying tibial spine heights (Table 2) and ensuring that the meniscus conformed to the tibial cartilage. The generation of the parametric model involved five steps:

**Table 2:**
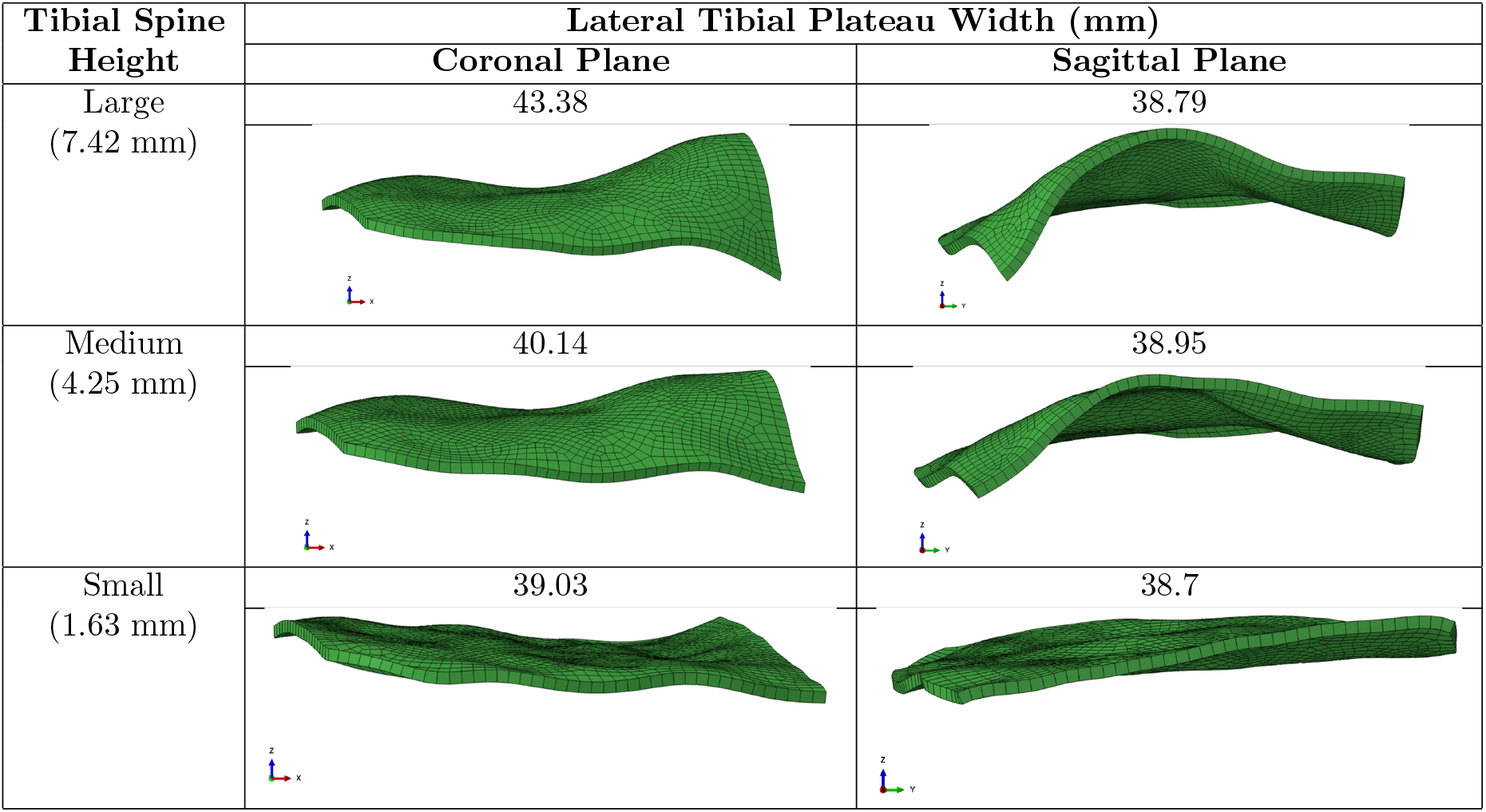
Geometrical parameters of the three finite element models of the lateral tibial plateau.

| Tibial Spine Height | Lateral Tibial Plateau Width (mm) |  |
| --- | --- | --- |
|  | Coronal Plane | Sagittal Plane |
| Large (7.42 mm) | 43.38 | 38.79 |
| Medium (4.25 mm) | 40.14 | 38.95 |
| Small (1.63 mm) | 39.03 | 38.7 |

- Cartilage Partitioning and Cross-Section Extraction. The geometry of tibial and femoral cartilage was obtained from scans and traced in SOLIDWORKS. To achieve a conformal meniscus model, the cartilage was partitioned into strips, with a spacing of 4 mm (Fig. 1c). This value was chosen arbitrarily to retain geometric fidelity while minimising the number of strips for practical modifications.
- Spline-based Top Surface Tracing. The top surface of each cartilage strip was traced using the spline tool in SOLIDWORKS. By defining the spline with a range of points, any adjustment to individual points updated the geometrical model, providing real-time control over cross-sections.
- Uniform Thickness Generation. The traced spline (Fig. 1d) was offset to create a controllable uniform thickness throughout the model. This feature ensures accurate representations of cartilage thickness variations, critical in biomechanical analyses.
- Meniscus Bottom Face Generation. The profile of the tibial cartilage’s top surface (the tibial cartilage component shown in yellow in Fig. 1e) served as a reference to generate the bottom face of the meniscus model. Geometrical parameters, namely width, length, and height, were estimated from scans, ensuring anatomical accuracy.
- Modeling Meniscus Structural Arrangements. Considering the depth-dependent material properties of menisci as observed in [22], the meniscus model was divided into three distinct regions: two superficial layers and a thicker mid-layer. The structural arrangement was achieved by offsetting the bottom and top faces of the meniscus model and then using the resulting faces to split the model. The layer depth was modified by altering one dimension, facilitating the creation of models with varying thicknesses (Fig. 1g).

### 2.3. FE model generation of knee joint lateral compartments

The 3D geometries of the lateral compartment of the knee joints (Fig. 1e) were imported into ABAQUS 2022 (Dassault Systèmes), generating FE models of the lateral condyle (Fig. 1f). Three-dimensional linear solid elements were used to mesh meniscus with C3D4 (4-node linear tetrahedron), and tibia and femur cartilage with C3D8RH (8-node linear brick, reduced integration with hourglass control, hybrid with constant pressure). Three FE models with varying tibial spine heights were generated to study the sensitivity of such parameters on load distribution and meniscus kinematics. The spine heights and tibial plateau width in the FE models Table 2 were within the values estimated from the arthrograms Table 1. To report results in this paper, we categorised the three FE models into large, medium and small tibial spine heights representing the FE models with 7.42, 4.25 and 1.63 mm tibial spine heights, respectively.

The meniscus roots (ligaments) were simulated using a series of linear elastic spring elements, with stiffness of 1 MPa (four spring elements at each end of the meniscus with k=0.25 N/mm) [23]. The anterior springs, representing the anterior root, connected the meniscus’s anterior end to a fully constrained reference point offset from the anterior meniscus end. The posterior springs, representing the posterior root, connected the meniscus’s posterior end to another fully constrained reference point offset from the posterior meniscus end. Spring elements allow meniscus root behaviours to be readily modified independent of 3D geometry.

The contact interaction between the articulating surfaces was defined as surface-to-surface contact, using penalty friction formulation and a friction coefficient value of 0.05. The outer surfaces of the articular cartilage were kinematically coupled to reference points for the application of external loads and boundary conditions. The tibia cartilage reference point was constrained in all directions, and the femur cartilage reference point had one degree of freedom allowing for vertical movements to apply an axial load of 1000 N (approximately 2-3 times an average healthy body weight [24]).

Cartilage and meniscus were modelled as neo-Hookean solids. Cartilage was assumed to be incompressible (C10 = 1.667, D1 = 0) while the meniscus was divided into three layers [25], following the experimental results collected in [22, 26, 27, 28]. The two outer layers were approximately 0.2 mm thick, stiffer and nearly incompressible (C10 = 2.614, D1 = 0.008). The internal layer was approximately 9.3 mm, softer and more compressible (C10 = 2.256, D1 = 0.205). Simulations were performed using ABAQUS static general solver using non-linear geometry and took approximately 30 minutes on two cores.

## 3. Results

### 3.1. Meniscus kinematics and root strains

Changes in the height of the tibial spine had a notable effect on the meniscus movement in the FE models Fig. 2. A reduction in the spine height, when comparing Large FE (Fig. 2a) to Medium FE (Fig. 2b), and Small FE (Fig. 2c), illustrated a corresponding increase in total meniscus displacement. The meniscus displacement increased by approximately 1.7 and 3.1 folds radially (mediolateral direction) and by approximately 1.1 and 2.9 folds circumferentially (anteroposterior direction) when the tibial spine height decreased from large to medium and then medium to small, respectively (Table 3). In other words, when the tibial spine height was reduced in the FE model by approximately 6 mm from large to small, the maximum displacement was increased by approximately 421% radially and 216% circumferentially, showing 204% larger changes in the radial direction. This indicates that radial displacement was more sensitive to changes in the spine height than circumferential displacement. However, the maximum displacements were lower in the radial than in the circumferential directions for all the FE models (Fig.2 d).

**Table 3:** The maximum displacements of the meniscus in the radial and circumferential directions, along with the root strains, were measured in three finite element models representing varying tibial spine heights.

| Tibial Spine Height | Maximum Meniscus Displacement (mm) |  | Total Maximum Principal Strain in Meniscus Roots (mm/mm) |
| --- | --- | --- | --- |
|  | Radial | Circumferential |  |
| **Large (7.42 mm)** | 1.56 | 2.95 | 5.08 (anterior root: 3.89, posterior root: 1.18) |
| **Medium (4.25 mm)** | 2.61 | 3.19 | 11.25 (anterior root: 6.77, posterior root: 4.48) |
| **Small (1.63 mm)** | 8.12 | 9.33 | 44.11 (anterior root: 21.92, posterior root: 22.19) |

**Figure 2:**
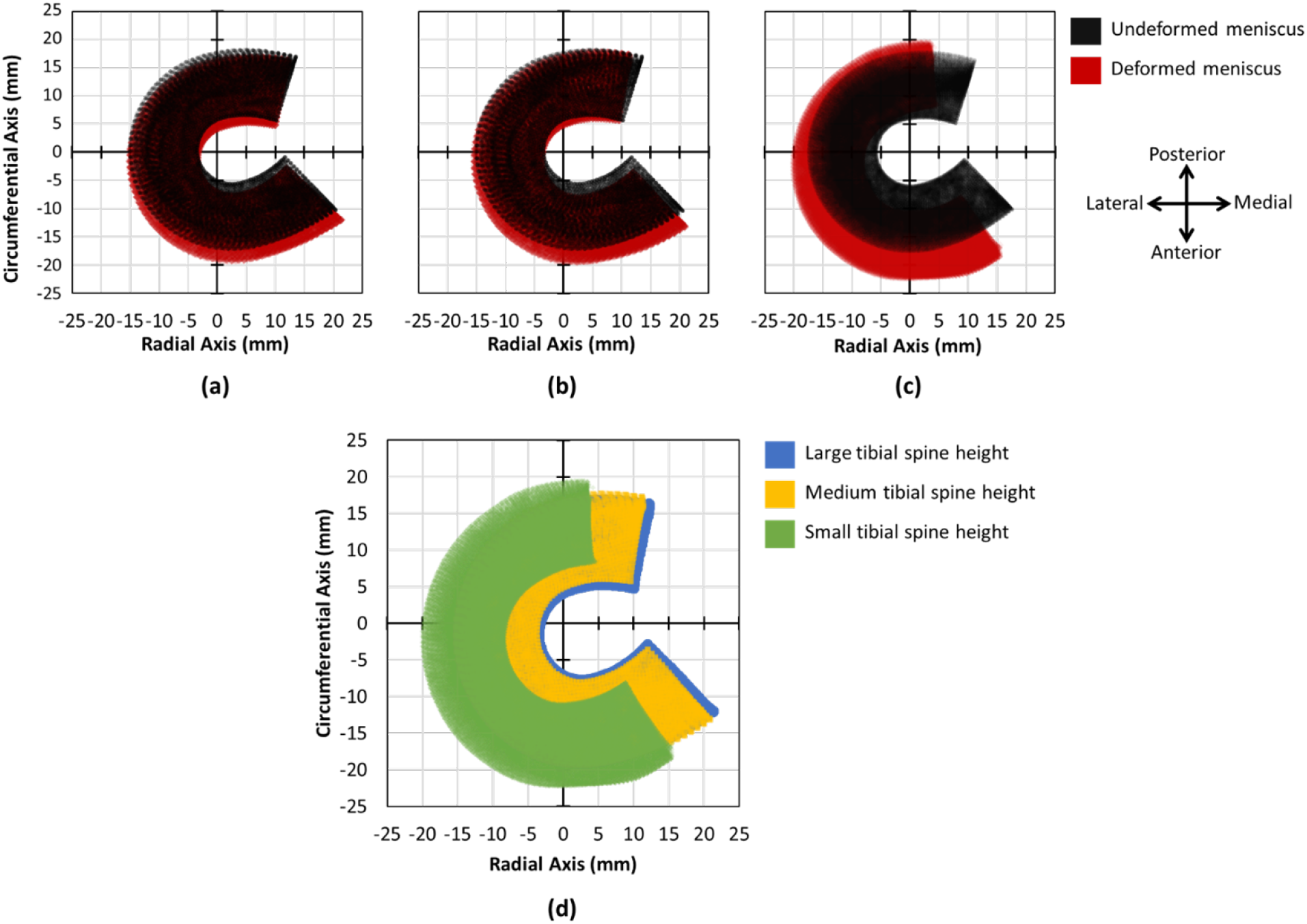
Comparisons of deformed (red) and undeformed (black) menisci in the radial and circumferential axes for the three knee joint models with (a) large, (b) medium and (c) small tibial spine heights. (d) Overlap of the deformed menisci illustrating the increase in circumferential and radial displacement with reducing tibial spine heights, meniscus in the high tibial spine is represented in blue, medium is yellow, and small is green).

The meniscus root strains increased with decreasing tibial spine height (Fig. 3). The root strains increased by approximately 2.2 and 3.9 folds when the tibial spine height decreased from large to medium and then medium to small (Table 3). The meniscus root strain was the largest (approximately 44 mm/mm) in the FE model with the smallest spine height, and the strain was evenly distributed between the anterior and posterior roots indicating symmetrical behaviour in the meniscus. With increased tibial spine height, the meniscus shape becomes increasingly unsymmetrical. The unsymmetrical shape led to an uneven strain in the roots of the FE models, especially in the medium and large models. The FE model with the highest spine height exhibited the largest uneven root strain, with the anterior root straining approximately three folds higher than the posterior root.

**Figure 3:**
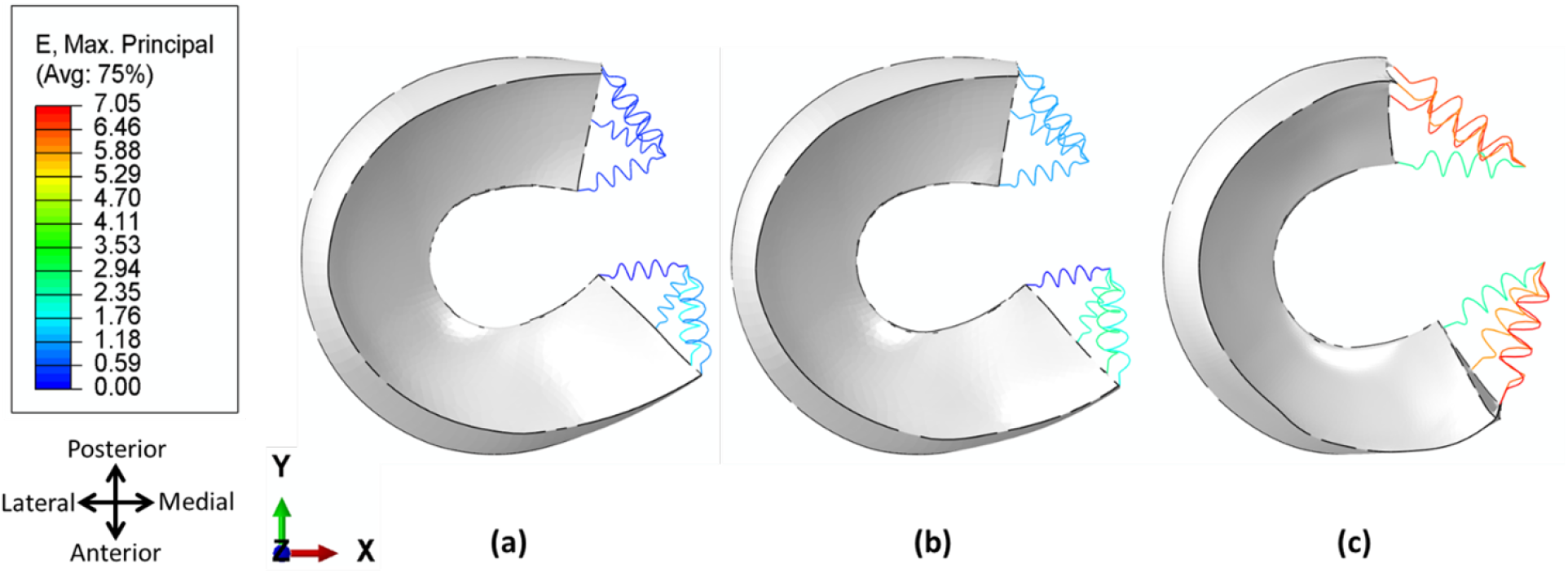
Maximum principal strains in meniscus roots represented by the spring elements in the three knee joint models with (a) high, (b) medium and (c) small tibial spine heights.

### 3.2. Contact mechanics in articulating surfaces

The contact mechanics (i.e., contact area and pressure) in the articulating surfaces were affected by changing tibial spine height (Table 4). The total contact area has increased while contact pressure was decreased with decreasing tibial spine height. The distribution of contact area between tibial cartilage on femoral cartilage (CoC) and meniscus on tibial cartilage (MoC) did not change noticeably. For example, the small spine height model predicted a total contact area of approximately 144 *mm*^2^, of which 84% was CoC and 16% was MoC. Similarly, the large spine height model showed a total contact area of approximately 78 *mm*^2^, of which 85% was CoC and 15% was MoC.

**Table 4:** Total contact area (mm^2^), force (N), and pressure (MPa) in femur cartilage on tibia cartilage (CoC) and meniscus on tibia cartilage (MoC) measured in the three finite element models with varying spinal height (small, medium and large).

| Tibial Spine Height | Total Contact Area ($\text{mm}^2$) | | Total Contact Force (N) | | Contact Pressure (MPa) | |
| --- | --- | --- | --- | --- | --- | --- |
|  | CoC | MoC | CoC | MoC | CoC | MoC |
| Large (7.42 mm) | 66.586 | 11.855 | 996.988 | 3.612 | 14.973 | 0.305 |
| Medium (4.25 mm) | 74.597 | 10.468 | 996.592 | 8.964 | 13.360 | 0.856 |
| Small (1.63 mm) | 121.281 | 22.918 | 962.442 | 37.382 | 7.936 | 1.631 |

The CoC surfaces showed a considerably higher contact force in all three FE models than the MoC surfaces. This suggests that the articulating cartilage plays a vital role in supporting at least 96% of the total load, as shown in Table 3. The load carried by the meniscus increased from approximately 4 to 9 and 37 N with decreasing tibial spine height from large to medium and then to small.

The mean contact pressure, calculated from the contact area and force, decreased in the CoC and increased in the MoC with decreasing tibial spine height (Table 3). For example, the contact pressure decreased by approximately 47% in the CoC and increased by 435% when tibial spine height decreased from large to small. The position of the contact pressure map on the articular cartilage was closer to the knee joint centre in the Large FE (Fig. 4a) model and has shifted laterally in the Medium (Fig. 4b) and Small (Fig. 4c) FE models. The contact pressures on the meniscus in the Large (Fig. 4d) and Medium (Fig. 4e) FE models were at the outer edge and horns of the meniscus. In contrast, contact pressure in the Small FE model (Fig. 4f) was more evenly distributed across the meniscus body.

**Figure 4:**
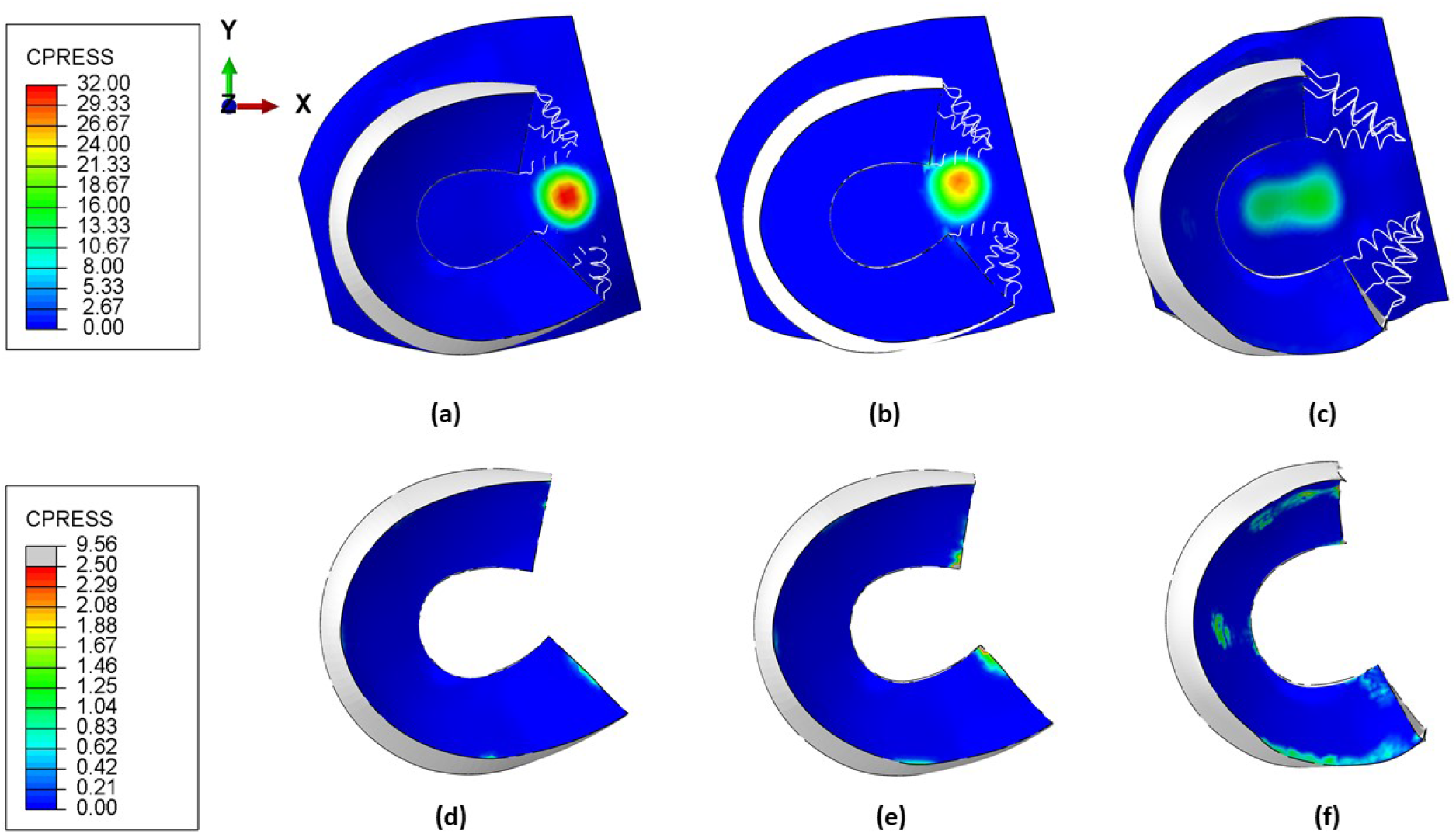
Contact pressure (MPa) maps in cartilage and meniscus for the three finite element models with (a and d) large, (b and e) medium, and (c and f) small tibial spine heights.

### 3.3. Meniscus stress and strain

The meniscus, comprised of two external layers and one internal layer, exhibited varying stress and strain patterns based on the height of the tibial spine (Fig. 5). Decreasing tibial spine heights increased stresses (Fig. 5) and logarithmic strains (6) in the meniscus structure. Meniscus in the FE model with the small tibial spine height experienced the highest stresses (Fig. 5c) and strains (6c) compared to the other two FE models. In the FE models, the outer layers of the meniscus showed the highest von Mises stress, with stress levels reaching approximately 0.5, 1.2, and 4 MPa for the large (Fig. 5a1 and a2), medium (Fig. 5b1 and b2), and small spine heights (Fig. 5c), respectively.

**Figure 5:**
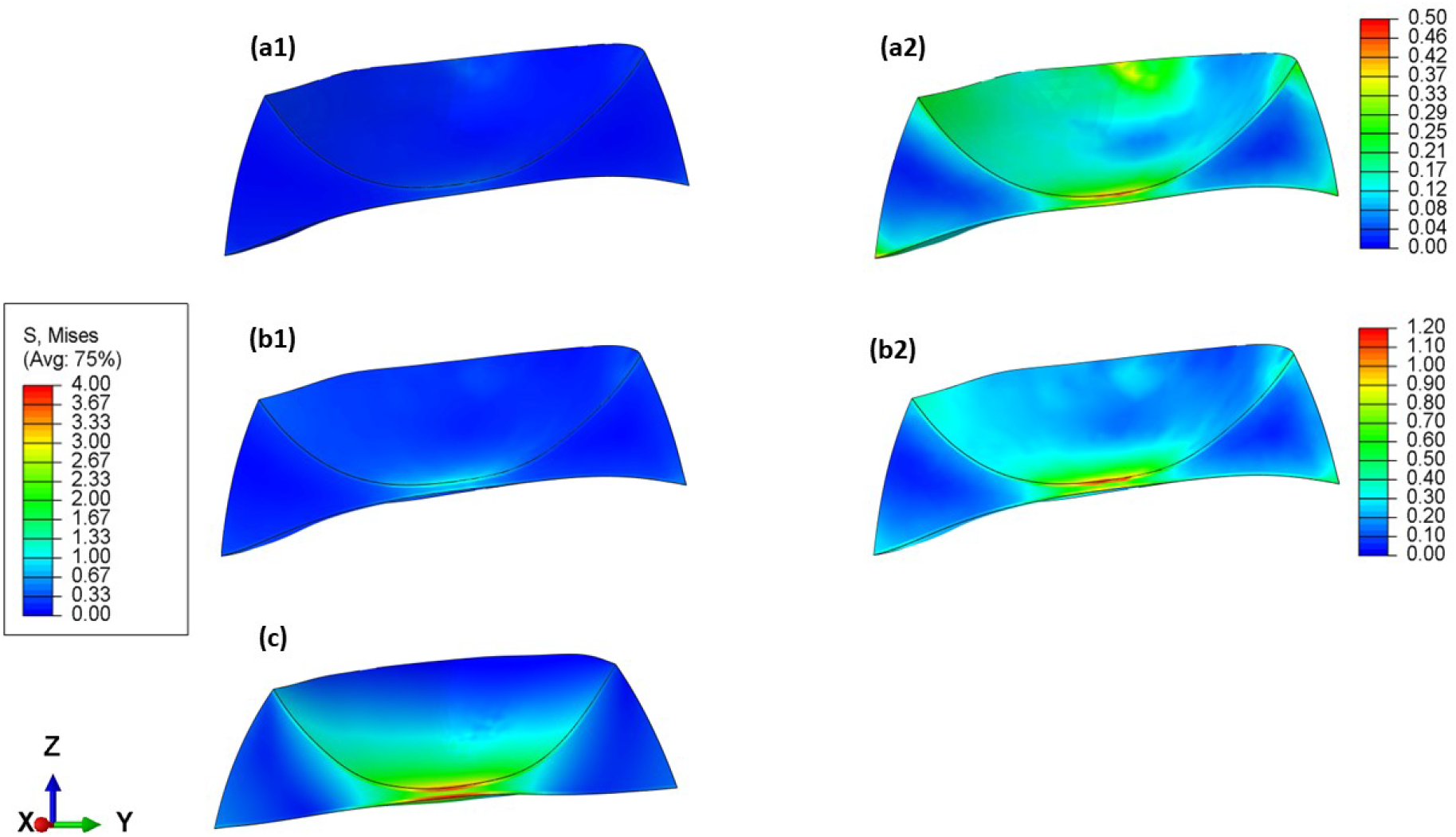
Von Mises stress(MPa) distribution across the three layers of the meniscus in the three finite element models with (a1 and a2) large, (b1 and b2) medium and (c) small tibial spine heights.

The logarithmic strains followed similar patterns to the stress distribution across the outer layers of the meniscus in the three FE models. For example, in the outer layers of the meniscus, the maximum logarithmic strains due to compressive load reached approximately 0.008, 0.023, and 0.138 mm/mm for the large (Fig. 6a1 and a2), medium (Fig. 6b1 and b2), and small spine heights (Fig. 6c), respectively. The inner layer of the meniscus shows strains due to tension present in the circumferential boundaries of the meniscus. The FE model with the lowest tibial spine height showed the highest tension-induced strain, reaching approximately 0.035 mm/mm (Fig. 6c). Conversely, the FE models with medium and large spine heights had strain values ranging between 0.010-0.018 mm, as depicted in Figures 6a2 and b2.

**Figure 6:**
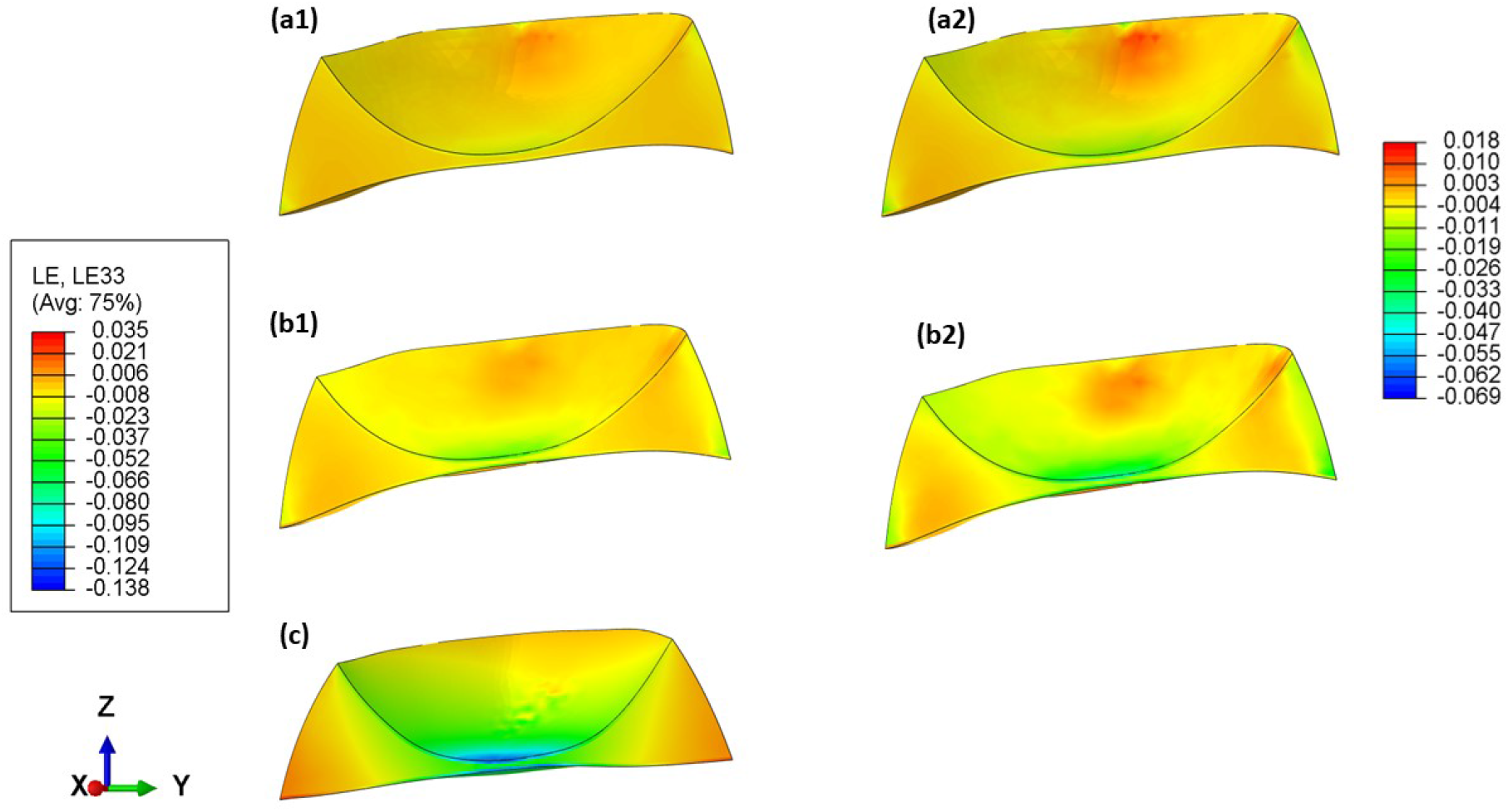
Logarithmic strains (mm/mm) across the three layers of the meniscus in the three finite element models with (a1 and a2) large, (b1 and b2) medium and (c) small tibial spine heights.

## 4. Discussion

A new framework has been introduced for creating patient-specific knee joint finite element models using arthrograms, eliminating the need for the time-consuming segmentation process typically used to convert medical images into 3D geometrical models. This study illustrated that creating models based on patient data using parametric methods instead of the typical image segmentation approach leads to improved simulation times, especially with respect to model convergence and enhanced solution quality at the contact surfaces. These models are highly adaptable for automation, allowing for the rapid production of comparable models for stochastic studies. The models could also be integrated into more comprehensive skeletal FE models, facilitating the simulation of dynamic loading cycles. The accuracy of defining and assigning effective mesh densities to the thin layers present in knee joint structures is a significant advantage of the framework described in this study. For example, parametric modelling provides a precise approach to defining the outer layers of a knee joint meniscus, allowing for optimal meshing and improved simulation. Here we tested the framework by generating three FE models of a single knee joint compartment. These models had different tibial spine heights categorised into small, medium and large based on the range measured from arthrograms. The result showed that the framework produces FE models capable of rapid numerical convergence and sensitive to changing key geometrical parameters (i.e., tibial cartilage morphology) important for meniscus kinematics and contact mechanics. The FE models showed that reducing the height of the tibial spine led to increased meniscus displacement, meniscus root strains, meniscus contact pressure, stresses, and compressive strains. At the same time, there was a decrease in contact pressure on the tibial cartilage. As per the models, a decrease in the height of the tibial spine also caused the contact area between the articulating surfaces to shift laterally away from the centre of the joint.

To our knowledge, no study in the literature described a framework for generating a parametric knee model with a particular focus on the knee joint meniscus. Most previous studies focused on generating subject-specific knee joint finite element models derived from clinical image segmentation [29], [30],[31], [32].

Li et al. used a whole knee joint FE model to study the influence of meniscus degenerative lesions on the change of knee biomechanics when applying a vertical compressive load of 1150 N [31]. The method for calculating meniscus extrusion in their FE models remains unclear, whether it was solely radial or a combination of radial and circumferential displacement. Nonetheless, they reported approximately 2.6 mm lateral meniscus extrusion in a healthy knee, aligning with our large and medium FE model observations (Fig. 2). An experimental study examined meniscus movement in the anterior and posterior directions based on different knee joint flexion. According to the study, the meniscus displayed asymmetrical behaviour, with the anterior horn moving by a maximum of 13 mm and the posterior horn moving by a maximum of 10 mm [33]. Our FE model captured the unsymmetrical behaviour as observed in the meniscus root strains, and the FE model with the highest spine height exhibited the largest uneven root strain, with the anterior root straining approximately three folds higher than the posterior root (Fig. 3 and Table 3).

Limited studies focused on the function of meniscus roots, with most of the previous literature on FE modelling knee joints describing the roots as a continuation of the meniscus tissue [30]. Researchers experimentally determined the hyperelastic strain behaviour of meniscus roots and have reported failure strains in lateral menisci roots ranging from 0.4 to 0.94 mm/mm and ultimate elongations ranging from 8 to 10 mm [34], [35]. Our study described the meniscus roots as linear elastic springs to achieve rapid numerical convergence. For this reason, the maximum principal root strains ranging from 3 to 22 mm/mm were higher than the experimental data reported in the previous literature. However, the elongations of the meniscus roots observed from maximum meniscus displacements (Table 3) were below the ultimate root elongations, improving our confidence in our models [34].

The FE modelling literature, using static simulations, predicted higher contact areas and lower contact pressures than the results reported using models from our framework. For example, Bao et al. applied an axial load of 1000 N on a whole knee FE model and reported the contact area for CoC and MoC to be approximately 260 mm^2^ and 240 mm^2^, and peak pressure for CoC and CoM be approximately 6 and 3 MPa, respectively [32]. In a human cadaveric experiment, researchers used Tekscans and applied an axial load of 1800 N to determine the contact area and pressure between articulating surfaces in the knee joint. The results showed that the combined CoC and MoC had a contact area of 570 mm^2^ and a pressure of 5 MPa [36]. The observed discrepancies between the current and previous studies are likely a consequence of the incomplete alignment and conformity of femoral cartilage with the top surface of the meniscus, leading to lower contact areas and higher contact pressures. The contact mechanics predicted in the current study were more aligned with the results described by Mononen et al., who reported a contact pressure range of 10 to 12 MPa during dynamic loading [37]. It can be difficult to compare our generic and simplified models with subject-specific FE models, as highlighted by Cooper et al.[20]. In their review, Cooper et al. described the challenges in knee modelling and stressed the importance of developing generic models, such as the ones generated by our framework, for sensitivity testing and parametric analysis to gain a more comprehensive understanding of patterns in various demographics [20].

Luczkiewicz et al. used the Open Knee model to study the influence of geometrical changes in the distribution of stresses with an emphasis on meniscus extrusion [29]. They modelled the meniscus as a continuous and uniform structure with linear elastic and transversely isotropic material properties. Their approach for modelling the meniscus and other knee components was similar to previous FE modelling studies [38]. However, unlike our proposed framework, these models do not fully capture the structural organisation of the meniscus, such as the three layers and layer-specific material properties [22]. It is possible that the difficulty in meshing and simulating thin solid layers arises from the construction of 3D geometries through segmentation processes, as these can frequently produce surfaces that are rough and non-uniform. The FE models predicted region-specific stresses and strain values across the cross-section of the meniscus Figs. 5 and 6. The FE models demonstrated a decrease in von Mises stresses with increasing the tibial spine height, indicating the sensitivity of the models to changing geometrical parameters. The outer layer of the meniscus exhibited the highest von Mises stress of 4 MPa as they bear the load and resist crack propagation, which falls within the stress range reported by Mononen et al. [37]. Despite using a single and continuous material block for modelling the meniscus, they reported maximum principal stress values ranging from 3 to 10 MPa [37]. Unlike previous studies [37], [31], the layered meniscus models demonstrated negative strains (due to compressive stress) and positive strains (due to tensile stress) around the meniscus circumference (6), hence better representing in vivo meniscus behaviour by demonstrating the effect of hoop stress [39].

Certain limitations exist within this study, as with any finite element modelling approach. However, the primary limitation is ensuring the model is valid and produces reliable results. The framework creates generic FE models with only a few patient parameters (such as bone geometries) rather than subject-specific 3D geometry; therefore, validating the generic models fully [20] is challenging. However, results were compared against cadaveric experimental and computational studies to increase confidence in the FE models. One of the remaining challenges in the current framework is ensuring precise bone alignment to achieve optimal joint congruence. Additionally, the FE models had uniform articular cartilage thickness and material properties. However, studies have emphasised the importance of non-uniform cartilage thickness and material properties [40], [41]. A significant correlation exists between localised contact pressure and material properties [40]. For example, cartilage is expected to be thicker in areas under higher loads. However, the framework can be adapted to capture varying cartilage thickness at the stage where the 3D geometry of the knee is constructed (Fig. 1d), hence expanding the purpose of the framework for research directly related to articular cartilage.

In conclusion, this study has developed a framework for generating simplified parametric FE models to yield reliable results in a rapid timescale. This framework allows for automated model generation, thus expanding the ability to investigate the impact of morphological (geometry and material) modifications. This approach offers a more rapid solution than the conventional method of relying on a single subject-specific model, especially for obtaining a more comprehensive understanding of patterns in various populations or aiding in surgical decisions that necessitate a quick turnaround. Our future work will expand on the framework’s capabilities by incorporating additional parametric adaptability, including articular cartilage thickness, or including other knee joint soft tissues, which could promote using our rapid computational simulation techniques for clinical decisions.

## Acknowledgement

O.B would like to acknowledge the European Union’s Horizon 2020 -EU.1.3.2. - Nurturing excellence by means of cross-border and cross-sector mobility under the Marie Sklodowska-Curie individual fellowship MSCA-IF-2017, MetaBioMec, Grant agreement ID: 796405.

## References

[1] L. Blankevoort, J.-H. Kuiper, R. Huiskes, H. J. Grootenboer, Articular contact in a three-dimensional model of the knee, Journal of biomechanics 24 (11) (1991) 1019–1031.

[2] V. C. Mow, S. C. Kuei, W. M. Lai, C. G. Armstrong, Biphasic creep and stress relaxation of articular cartilage in compression: theory and experiments, Journal of biomechanical engineering 102 (1) (1980) 73–84.

[3] T. L. Donahue, M. L. Hull, M. S. Rashid, An experimental and numerical analysis of the contact stresses in the meniscus following partial meniscectomy, Journal of biomechanical engineering 124 (2) (2002) 138–145.

[4] M. E. Mononen, J. S. Jurvelin, R. K. Korhonen, M. Hauta-Kasari, Three-dimensional ultrasound imaging provides accurate full-thickness cartilage thickness measurements in porcine knee cartilage, Osteoarthritis and cartilage 20 (7) (2012) 686–692.

[5] A. E. Anderson, C. L. Peters, B. D. Tuttle, J. A. Weiss, Subject-specific finite element model of the pelvis: development, validation, and sensitivity studies, Journal of biomechanical engineering 127 (3) (2005) 364–373.

[6] I. Kutzner, B. Heinlein, F. Graichen, A. Bender, A. Rohlmann, A. M. Halder, …, G. Bergmann, Loading of the knee joint during activities of daily living measured in vivo in five subjects, Journal of biomechanics 43 (11) (2010) 2164–2173.

[7] E. G. Sutter, M. R. Widmyer, G. M. Utturkar, C. E. Spritzer, W. E. Garrett Jr, L. E. DeFrate, In vivo measurement of localized tibiofemoral cartilage strains in response to dynamic activity, American Journal of Sports Medicine 43 (2) (2015) 370–376.

[8] G. Li, L. E. DeFrate, Zonal variation in the material properties of the cartilage of the knee joint, Journal of biomechanics 37 (5) (2004) 639–644.

[9] P. Hauseux, J. S. Hale, S. P. A. Bordas, Calculating the Malliavin derivative of some stochastic mechanics problems, PLOS ONE 12 (12) (12 2017).

[10] N. Shrive, J. O’connor, J. Goodfellow, Load-bearing in the knee joint., Clinical Orthopaedics and Related Research (1976-2007) 131 (1978) 279–287.

[11] G. Li, L. E. DeFrate, H. Sun, T. J. Gill, H. E. Rubash, In vivo elongation of the anterior cruciate ligament and posterior cruciate ligament during knee flexion, American Journal of Sports Medicine 33 (7) (2005) 907–914.

[12] A. M. Kiapour, M. M. Murray, B. C. Fleming, Biomechanical outcomes of bridge-enhanced anterior cruciate ligament repair are influenced by sex in a preclinical model, Clinical Orthopaedics and Related Research 472 (8) (2014) 2396–2405.

[13] H. Kainz, K. Hajek, B. J. Schwaiger, D. Salaberger, D. Rieder, S. Nürnberger, …, A. Bauer, Mechanical analysis of individual knee joint structures—a step towards patient-specific finite element models, PLoS One 11 (3) (2016) e0152504.

[14] L. Esposito, Y. Yu, J. Li, C. Cosme, J. Xu, L. Yao, …, J. P. DeAngelis, Patient-specific anatomical knee joint model based on clinical imaging: A feasibility study, Journal of Orthopaedic Research 37 (3) (2019) 678–686.

[15] D. Mancini, G. Marchiori, F. Baruffaldi, V. Parenti-Castelli, Realistic 3d finite element model of the knee for the estimation of ligament properties, Computer Methods in Biomechanics and Biomedical Engineering 16 (11) (2013) 1173–1184.

[16] E. Elmukashfi, G. Marchiori, M. Berni, G. Cassiolas, N. F. Lopomo, H. Rappel, M. Girolami, O. Barrera, Chapter five -model selection and sensitivity analysis in the biomechanics of soft tissues: A case study on the human knee meniscus, Vol. 55 of Advances in Applied Mechanics, Elsevier, 2022, pp. 425–511. doi:10.1016/bs.aams.2022.05.001. URL https://www.sciencedirect.com/science/article/pii/S0065215622000023

[17] E. G. Sutter, J. D. Widmer, Y. Ji, R. M. Degen, K. W. Nha, W. R. Fitz, V. Juras, K. P. Pritzker, G. E. Gold, B. A. Hargreaves, et al., Semiautomated 3d magnetic resonance imaging segmentation of the knee extensor mechanism, Knee Surgery, Sports Traumatology, Arthroscopy 26 (1) (2018) 63–69.

[18] Y. Chen, X. Zhao, H. Zhong, Z. Liao, L. Yang, S. Li, H. Lu, Y. Hu, Automatic segmentation of knee bones and cartilages combining statistical shape models and convolutional neural networks: data from the osteoarthritis initiative, Frontiers in Bioengineering and Biotechnology 7 (2019) 171.

[19] M. F. Rai, E. J. Schmidt, D. R. McAllister, M. R. Schmitz, K. T. Vann, R. E. Stroud, F. Guilak, The meniscus: basic science, mechanical properties, and tissue engineering strategies, Journal of Biomechanical Engineering 139 (2) (2017) 021010.

[20] R. J. Cooper, R. K. Wilcox, A. C. Jones, Finite element models of the tibiofemoral joint: A review of validation approaches and modelling challenges, Medical Engineering and Physics 74 (2019) 1–12. doi:10.1016/j.medengphy.2019.08.002. URL https://www.sciencedirect.com/science/article/pii/S1350453319301572

[21] K. Halonen, M. Mononen, J. Jurvelin, J. Töyräs, J. Salo, R. Korhonen, Deformation of articular cartilage during static loading of a knee joint – experimental and finite element analysis, Journal of Biomechanics 47 (10) (2014) 2467–2474. doi:10.1016/j.jbiomech.2014.04.013. URL https://www.sciencedirect.com/science/article/pii/S0021929014002346

[22] J. Maritz, G. Agustoni, K. Dragnevski, S. P. A. Bordas, O. Barrera, The functionally grading elastic and viscoelastic properties of the body region of the knee meniscus, Annals of Biomedical Engineering (2021) 1–9.

[23] G. B. B. K. T. A. R. R. R. C. E. Peters, A. E., Ligament mechanics of ageing and osteoarthritic human knees, Frontiers in bioengineering and biotechnology 10 (2022) 954837. doi:10.3389/fbioe.2022.954837.

[24] H. J. S. Z. J. A. Knarr, B. A., Change in knee contact force with simulated change in body weight, Computer methods in biomechanics and biomedical engineering 19 (3) (2016) 320–323. doi:10.1080/10255842.2015.1018193.

[25] G. Agustoni, J. Maritz, J. Kennedy, F. P. Bonomo, S. P. A. Bordas, O. Barrera, High resolution micro-computed tomography reveals a network of collagen channels in the body region of the knee meniscus, Annals of Biomedical Engineering (2021) 1–9.

[26] F. P. Bonomo, J. J. S. Gregory, O. Barrera, A procedure for slicing and characterizing soft heterogeneous and irregular-shaped tissue, Materials Today: Proceedings 33 (2020).

[27] J. Maritz, F. Murphy, K. Dragnevski, O. Barrera, Development and optimisation of micromechanical testing techniques to study the properties of meniscal tissue, Materials Today: Proceedings (2020).

[28] V. Vetri, K. Dragnevski, M. Tkaczyk, M. Zingales, G. Marchiori, N. F. Lopomo, S. Zaffagnini, A. Bondi, J. A. Kennedy, D. W. Murray, O. Barrera, Advanced microscopy analysis of the micro-nanoscale architecture of human menisci, Scientific Reports 9 (1) (2019) 1–13.

[29] D. K. W. W. C. J. Z. W. Luczkiewicz, P., Influence of meniscus shape in the cross sectional plane on the knee contact mechanics, Journal of biomechanics 48 (8) (2015) 1356–1363. doi:10.1016/j.jbiomech.2015.03.002.

[30] E. Peña, M. M. A. D. M. Calvo B., A three-dimensional finite element analysis of the combined behavior of ligaments and menisci in the healthy human knee joint., Journal of biomechanics 39 (9) (2006) 1686–1701. doi:10.1016/j.jbiomech.2005.04.030.

[31] Y. L. Z. K. Z. L. W. X. J. Q. Li, L., Three-dimensional finite-element analysis of aggravating medial meniscus tears on knee osteoarthritis, Journal of orthopaedic translation 20 (2019) 47–55. doi:10.1016/j.jot.2019.06.007.

[32] Z. D. G. H. G. G. S. Bao, H. R., The effect of complete radial lateral meniscus posterior root tear on the knee contact mechanics: a finite element analysis, Journal of orthopaedic science: official journal of the Japanese Orthopaedic Association 18 (2) (2013) 256–263. doi:10.1007/s00776-012-0334-5.

[33] T. F. L. F. F. H. D. S. F. Thompson, W. O., Tibial meniscal dynamics using three-dimensional reconstruction of magnetic resonance images, The American journal of sports medicine 19 (3) (1991) 210–216. doi:10.1177/036354659101900302.

[34] M. J. A. M. S. D. D. T. L. Villegas, D. F., Failure properties and strain distribution analysis of meniscal attachments, Journal of biomechanics 40 (12) (2007) 2655–2662. doi:10.1016/j.jbiomech.2007.01.015.

[35] H. M. L. R. M. M. J. C. R. Haut Donahue, T. L., How the stiffness of meniscal attachments and meniscal material properties affect tibio-femoral contact pressure computed using a validated finite element model of the human knee joint. journal of biomechanics, Journal of orthopaedic science: official journal of the Japanese Orthopaedic Association 36 (1) (2003) 19–34. doi:10.1016/s0021-9290(02)00305-6.

[36] . G.-D. J. Marzo, J. M., Effects of medial meniscus posterior horn avulsion and repair on tibiofemoral contact area and peak contact pressure with clinical implications, The American journal of sports medicine 37 (1) (2009) 124–129. doi:10.1177/0363546508323254.

[37] J. J. S. K. R. K. Mononen, M. E., Effects of radial tears and partial meniscectomy of lateral meniscus on the knee joint mechanics during the stance phase of the gait cycle–a 3d finite element study, Journal of orthopaedic research: official publication of the Orthopaedic Research Society 31 (8) (2013) 1208–1217. doi:10.1002/jor.22358.

[38] A. R. C. E. J. B. K. T. Peters, A. E., Tissue material properties and computational modelling of the human tibiofemoral joint: a critical review, PeerJ 6 (2018) e4298. doi:10.7717/peerj.4298.

[39] A. J. S. Fox, A. Bedi, S. A. Rodeo, The basic science of human knee menisci: Structure, composition, and function, Sports Health 4 (4) (2012) 340–351.

[40] S. C. R. Z. L. T. D. G. V. B. V. A. D. J. I. Van Rossom, S., Knee cartilage thickness, t1 and t2 relaxation time are related to articular cartilage loading in healthy adults, PloS one 12 (1) (2017) e0170002. doi:10.1371/journal.pone.0170002.

[41] G. M. D. E. Zanjani-Pour, S., Development of subject specific finite element models of the mouse knee joint for preclinical applications, Frontiers in bioengineering and biotechnology 8 (2020) 558815. doi:10.3389/fbioe.2020.558815.

